# Repeat Detector: versatile sizing of expanded tandem repeats and identification of interrupted alleles from targeted DNA sequencing

**DOI:** 10.1101/2022.03.08.483398

**Authors:** Alysha S. Taylor, Dinis Barros, Nastassia Gobet, Thierry Schuepbach, Branduff McAllister, Lorene Aeschbach, Emma L. Randall, Evgeniya Trofimenko, Eleanor R. Heuchan, Paula Barszcz, Marc Ciosi, Joanne Morgan, Nathaniel J. Hafford-Tear, Alice E. Davidson, Thomas H. Massey, Darren G. Monckton, Lesley Jones, REGISTRY Investigators of the European Huntington’s disease network, Ioannis Xenarios, Vincent Dion

## Abstract

Targeted DNA sequencing approaches will improve how the size of short tandem repeats is measured for diagnostic tests and pre-clinical studies. The expansion of these sequences causes dozens of disorders, with longer tracts generally leading to a more severe disease. Interrupted alleles are sometimes present within repeats and can alter disease manifestation. Determining repeat size mosaicism and identifying interruptions in targeted sequencing datasets remains a major challenge. This is in part because standard alignment tools are ill-suited for repetitive and unstable sequences. To address this, we have developed Repeat Detector (RD), a deterministic profile weighting algorithm for counting repeats in targeted sequencing data. We tested RD using blood-derived DNA samples from Huntington’s disease and Fuchs endothelial corneal dystrophy patients sequenced using either Illumina MiSeq or Pacific Biosciences single-molecule, real-time sequencing platforms. RD was highly accurate in determining repeat sizes of 609 blood-derived samples from Huntington’s disease individuals and did not require prior knowledge of the flanking sequences. Furthermore, RD can be used to identify alleles with interruptions and provide a measure of repeat instability within an individual. RD is therefore highly versatile and may find applications in the diagnosis of expanded repeat disorders and the development of novel therapies.

## Introduction

Huntington’s disease (HD) is one of the best studied members of a family of disorders caused by the expansion of short tandem repeats (1). It is characterised by neurodegeneration in the striatum and cortex, leading to chorea, cognitive decline, and premature death (2). The size of the inherited CAG repeat tract at the huntingtin (*HTT*) locus accounts for about 60% of the variability in the age at motor disease onset (3, 4), with longer repeats associated with earlier onset. Consequently, it is not possible to predict HD onset solely based on *HTT* repeat size, highlighting the importance of other factors contributing to disease pathology. One such factor is likely to be somatic expansion, or the ongoing expansion of expanded repeats in affected tissues throughout an individual’s lifetime (5). The contribution of somatic expansion to pathogenesis is highlighted by the number of genes implicated in repeat instability that also appear to modify age at disease onset (6, 7). It also follows that if ongoing somatic expansion contributes to disease phenotypes, gains or losses of interruptions within the repeat tract should lead to changes in the age at disease onset. We see that when repeats are interrupted, the repeat tract is stabilised and there correlates a later appearance of disease symptoms (6, 8). About 95% of HD chromosomes have a CAACAG motif immediately 3⍰ to the end of the CAG repeat tract, often referred to as an interruption (8). Alleles without this CAACAG interruption are associated with an earlier onset than predicted based on their repeat size (6, 8–11), whereas those with two CAACAG were either found to have no effect on the age at disease onset (6, 8) or were associated with a later onset (10).

Both somatic expansion and repeat interruptions also appear to influence disease outcome in other expanded repeat disorders (12). For example, somatic expansion is seen in affected tissues in myotonic dystrophy (13–16). Moreover, some of the genetic modifiers of HD implicated in repeat expansion may also modify disease onset in other repeat disorders (17). Interruptions in SCA1, SCA2, Fragile X syndrome, and myotonic dystrophy type 1 are associated with lower repeat instability, delayed symptom onset, and/or modified clinical manifestations (15, 16, 26–28, 18–25).

Interruptions are difficult to find using current PCR-based diagnostic tools (23, 29), and repeat instability is not currently measured in the clinic. The advent of high-throughput sequencing offers an opportunity to improve diagnosis by enhancing the accuracy of repeat sizing as well as the identification of interrupted alleles. Targeted sequencing of expanded repeats has been achieved with Illumina MiSeq (8, 10, 30), Pacific BioSciences (PacBio) Single-Molecule, Real Time (SMRT) (23, 26, 29–36), and Oxford Nanopore Technology MinION (32, 37–39). One of the remaining bottlenecks is the robustness of computational pipelines that can reliably determine repeat size and repeat interruptions at the single-molecule level in targeted sequencing datasets. Current algorithms (8, 40–43) all rely on the alignment of each read to a reference sequence. The presence of a highly variable tandem repeat can result in the rejection of read from the dataset, thereby introducing biases. The alignment step also limits the application of these algorithms to specific loci or genomes. Importantly, only one currently available algorithm allows for the unsupervised identification of novel interrupted alleles, the proprietary RepeatAnalysisTools by Pacific Biosciences, but it only works on data generated using the amplification-free library preparation for SMRT sequencing (29, 33). Here we present Repeat Detector (RD), a versatile algorithm that accurately counts expanded repeats in targeted sequencing datasets and can identify interrupted alleles. It works on datasets from multiple loci, sequencing platforms, and repeated motifs, making it widely applicable.

## Methods

### Cell culture and cell lines

The GFP(CAG)_x_ cell lines were cultured as described before (44, 45). The culture medium used was Gibco™ Dulbecco’s modified Eagle’s medium (DMEM) with GlutaMAX™, 10% foetal bovine serum (FBS), 100⍰U⍰ml^−1^ of penicillin/streptomycin, 15⍰μg⍰ml^−1^ blasticidine and 150⍰μg⍰ml^−1^ hygromycin. The HD lymphoblastoid cells (LBCs) or their DNA used for SMRT sequencing were obtained from the Coriell BioRepository (Table S1). The LBCs were grown in Gibco^™^ RPMI with GlutaMAX^™^ supplemented with 15% Gibco^™^ FBS (Thermo Fisher), and 1% penicillin-streptomycin. Both the LBCs and the GFP(CAG)_x_ cells were grown at 37⍰°C with 5% CO_2_ and tested negative for mycoplasma by Eurofins’ ‘Mycoplasmacheck’ service.

GFP(CAG)_91_ is identical to the previously characterised GFP(CAG)_101_ (45) but contained a contraction in the cultures used here. Similarly, GFP(CAG)_51_ had a one CAG expansion compared to when it was first derived (45) and GFP(CAG)_308_ had a repeat tract above 270 that we could not fully sequence with Sanger sequencing at the time. GFP(CAG)_15_, GFP(CAG)_51_, and GFP(CAG)_308_ are derived from GFP(CAG)_101_.

### Confirmation of interruption

We confirmed the presence of an interruption in GFP(CAG)_308_ by first amplifying the repeat region using primers oVIN-459 and oVIN-460 (for primer sequences see Table S2) and then Sanger sequencing using the same primers. The Sanger sequencing was done by GeneWiz.

### SMRT sequencing

The HD LBCs and GFP(CAG)_x_ datasets were generated by first isolating DNA using the Macherey-Nagel Nucleospin™ Tissue Mini kit. PCR products were generated from samples using barcoded primers as listed in Table S1 and Thermo™ Phusion II High Fidelity polymerase. To obtain sufficient quantities of PCR product to proceed with library preparation, multiple identical PCRs were pooled and purified using Macherey-Nagel™ Gel and PCR Clean-up kit columns. The library was generated using the SMRTbell Template Prep Kit (1.0-SPv3) according to manufacturer’s instructions. Samples to be sequenced on the same flowcell were combined in equimolar pools. We loaded between 10 and 12 pM. SMRT sequencing was done using a Sequel at Cardiff University School of Medicine. CCSs were generated from the resulting sequences and processed using SMRT Link.

### Participants

Human subjects were selected from the European Registry-HD study (46) (N=507) (https://www.enroll-hd.org/enrollhd_documents/2016-10-R1/registry-protocol-3.0.pdf) Ethical approval for Registry was obtained in each participating country. Participants gave written informed consent. Experiments described herein were conducted in accordance with the Declaration of Helsinki. Institutional ethical approval was gained from Cardiff University School of Medicine Research Ethics Committee (19/55). Subjects were selected as in (10).

### HD MiSeq dataset

A total of 652 DNA samples were sequenced, with the majority of these being immortalised lymphoblastoid (LBC) cell lines (N=547) and a smaller number of blood DNAs (N=49). These were sequenced using an ultra-high depth MiSeq sequencing methodology, described elsewhere (30, 47). Of note, the method includes a size selection step that biases towards longer alleles. 649 of the original 652 samples were successfully sequenced (>99%). Table S2 describes the numbers of each sample as well as the numbers of each DNA type that was successfully sequenced.

### FECD SMRT dataset

The FECD SMRT dataset is a amplification-free SMRT sequencing dataset from blood samples of FECD patients published previously (33).

### Repeat Detector

Repeat Detector source code and dependencies are available at: https://github.com/DionLab/RepeatDetector. To determine repeat sizes for GFP(CAG)_x_, FECD (33), HD MiSeq (10) and c9orf72 loci (37) datasets, unaligned reads were assessed using permissive and restrictive profiles with a repeat size range of [0-1000]. For each analysis, the -- with-revcomp option was enabled and data was output to a density plot (-o histogram option). Weighting scores for the permissive and restrictive parameters can be found in Fig. S1. Density plots obtained were graphed using GraphPad PRISM version 9.

### ScaleHD

The ScaleHD parameters were set as previously (10). For comparisons between ScaleHD and RD presented in Fig. 3B, Fig. S2, and Fig. S3, we used the total number of reads mapping to the *HTT* locus in the R1 FASTA files, regardless of the flanking sequences that sometimes differed between reads from the same sample. These are due to PCR and sequencing errors.

### Tandem-Genotypes

The FECD SMRT dataset (33) was aligned to GRCh38.18 accessed from the Genome Reference Consortium (48) using the LAST aligner (49), as per recommendation in (42). The reference sequence was soft-masked as per LAST aligner guidelines (https://github.com/mcfrith/last-rna) and sequences were aligned using default settings as described in the wiki. Aligned sequences were examined for the FECD repeat using Tandem-Genotypes recommended settings and modal repeat sizes were extracted from the output files.

## Results

### Repeat Detector

RD (Fig. 1) is based on the deterministic profile weighting algorithm, pfsearchV3.0, which was originally designed for protein motifs and domain detection (50, 51). It has been adapted to use circular profiles on DNA sequences. RD is not dependent on an alignment to a specific reference sequence. Instead, the user defines the repeated motif and the weighting parameters. RD then aligns the reads to a circular representation of the motif of interest. The weights of the profile give flexibility to adapt the alignment scoring to prior knowledge, for example about the idiosyncratic errors of a given sequencing platform or for the repeated motif of interest.

**Fig. 1:**
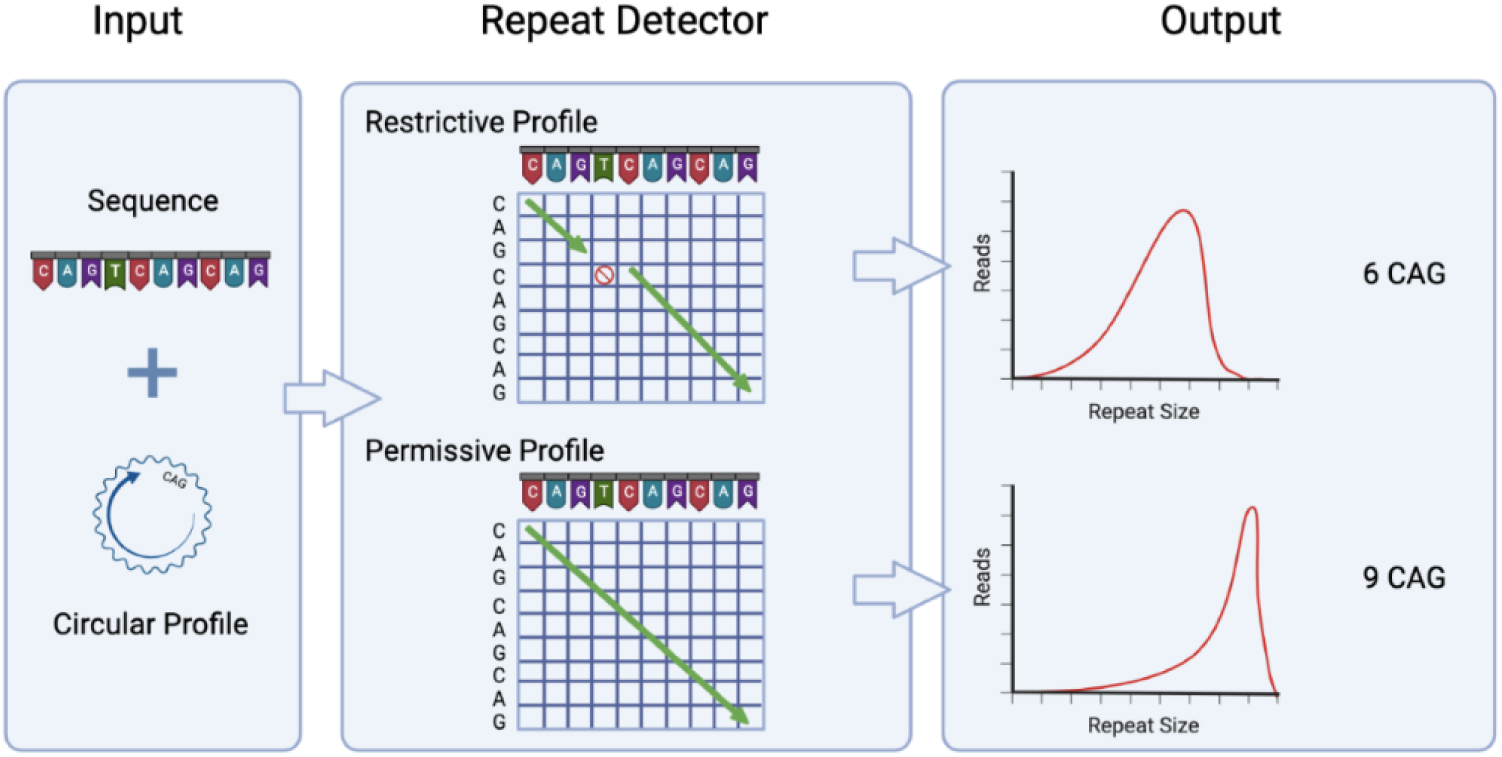
Repeat Detector flowchart. RD requires both the FASTA files of the DNA sequences and the circular profile of the repeating motif of interest as inputs. Using a substitution matrix, it calculates a score, taking into account matches, mismatches, gaps, and insertions. The repeat size with the largest score is deemed to be the correct one. There are two sets of parameters described in the methods. One is permissive and is lenient with non-matching nucleotides. The other is restrictive and stops counting when a mismatch, gap, or insertion is encountered. RD outputs the frequencies of repeat sizes, which are then presented as density plots.

### RD applied to two different loci over a wide range of repeat sizes

We first tested RD on two different datasets generated using SMRT sequencing and a standard PCR-based library preparation method. SMRT sequencing uses rolling circle replication chemistry that generates reads with multiple copies of the target sequence called subreads. A proprietary bioinformatics tool generates circular consensus sequences (CCSs) from subreads, improving base calling accuracy (52). Our first dataset consisted of CCSs from HEK293-derived cell lines with 15, 51, 91, and 308 CAG/CTG repeats inserted within a hemizygous ectopic GFP reporter on chromosome 12 (45, 53, 54). We refer to these cells as GFP(CAG)_x_, with x being the number of repeats. These lines are single-cell isolates derived from the previously characterised GFP(CAG)_101_ line (see methods and (45)). The second dataset was composed of 21 DNA samples and LBCs from HD individuals obtained from the Coriell BioRepository with repeats ranging from 15 to 750 units (Table S1). Taking both datasets together, we recovered the expected repeat sizes based on Coriell’s data or our prior work (55), except for one sample (Fig. 2ab). Only the sample with the longest repeat tract, GM14044, which we have shown to contain 750 repeats (55), returned a repeat size of 50 CAGs. By inspecting reads manually, we confirmed that the sequences in the FASTA files used by RD contained repeat sizes shorter than 750 repeats, suggesting that, rather than a specific problem with RD, there was a bias against longer repeats during PCR, loading of the SMRT flowcell, sequencing, and/or the generation of CCSs. These results are in line with recent findings suggesting that up to at least 550 CAG repeats can be sequenced using SMRT sequencing (29, 30).

**Fig. 2:**
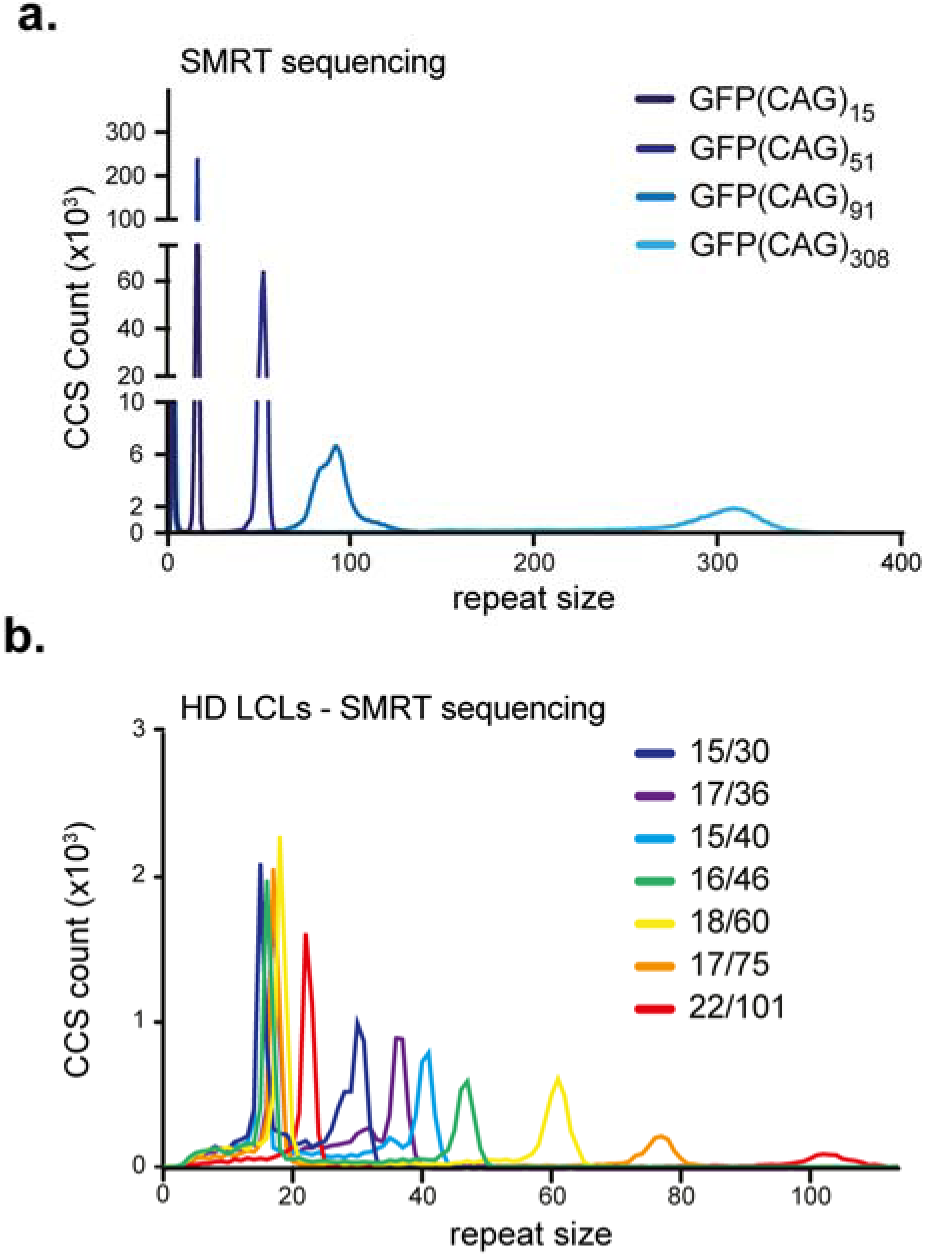
Repeat Detector applied to SMRT sequencing of an ectopic CAG/CTG repeat and at the HTT locus. A) RD-generated repeat size distribution from SMRT sequenced of ectopic CAG repeats in GFP(CAG)x cell lines using a PCR-based library preparation. B) Repeat size distribution of SMRT-sequenced samples of the HTT locus from HD-derived LBCs using a PCR-based library preparation. Only a selection of the 22 samples are shown for clarity. All samples are shown in Fig. S4. Read depth and mapping metrics for all datasets can be found in Table S4.

### RD is highly accurate on HD samples

We next sought to quantify the accuracy of RD in sizing clinically relevant samples. To do so, we took advantage of a previously sequenced set of 649 samples derived from 507 clinically manifesting HD individuals (10). This cohort included samples from 497 LBC lines, 49 blood samples sequenced twice, 47 LBC samples that were passaged extensively and an additional seven LBC samples from a single HD individual with a known repeat length, which ensured reproducibility (Table S3). For 42 individuals, there are data for both blood and LBCs. Hereafter, we refer to this dataset as the HD MiSeq dataset since it was generated using Illumina MiSeq technology (10). This dataset was originally analysed for modal repeat size and flanking sequences using ScaleHD (47). This algorithm uses a library containing over four thousand reference sequences with all known flanking sequences as well as repeat sizes between 1 and 200 CAGs. This created a robust benchmark against which we could evaluate RD for its ability to determine repeat size. Of the 649 samples, we analysed 609 with both algorithms, totalling 1218 germline alleles (Fig. 3a). For the shorter alleles, the modal repeat size was determined to be the same with both softwares (Fig. 3b). Of the longer alleles, 599 out of 609 (98.3%) had the same modal allele size (Fig. 3b). Of the remaining ten alleles, nine differed by one CAG and one allele by two (Fig. S2). One of these differences came from a homozygous individual with 2 alleles of 15 repeats. The script, downstream of RD, looks for the two most common allele sizes and thus determined erroneously that this sample had one allele with 15 repeats and one with 14 (Fig. S3a). The sample that differed most between ScaleHD and RD was an LBC sample derived from a confirmed HD individual. We had several samples from the same individual, yet ScaleHD determined this LBC sample to have two alleles with 19 repeats (Fig. S3b). RD, on the other hand, found one allele with 19 repeats and one with 42, in line with the other samples from this individual. The discrepancy was due to ScaleHD filtering out much of the reads containing the expanded allele. It is unclear why this occurred. RD does not rely on an alignment to the locus of interest and thus counted both alleles accurately (Fig. S3b). These data highlight the accuracy of RD and show that it is comparable to ScaleHD for the *HTT* locus.

**Fig. 3:**
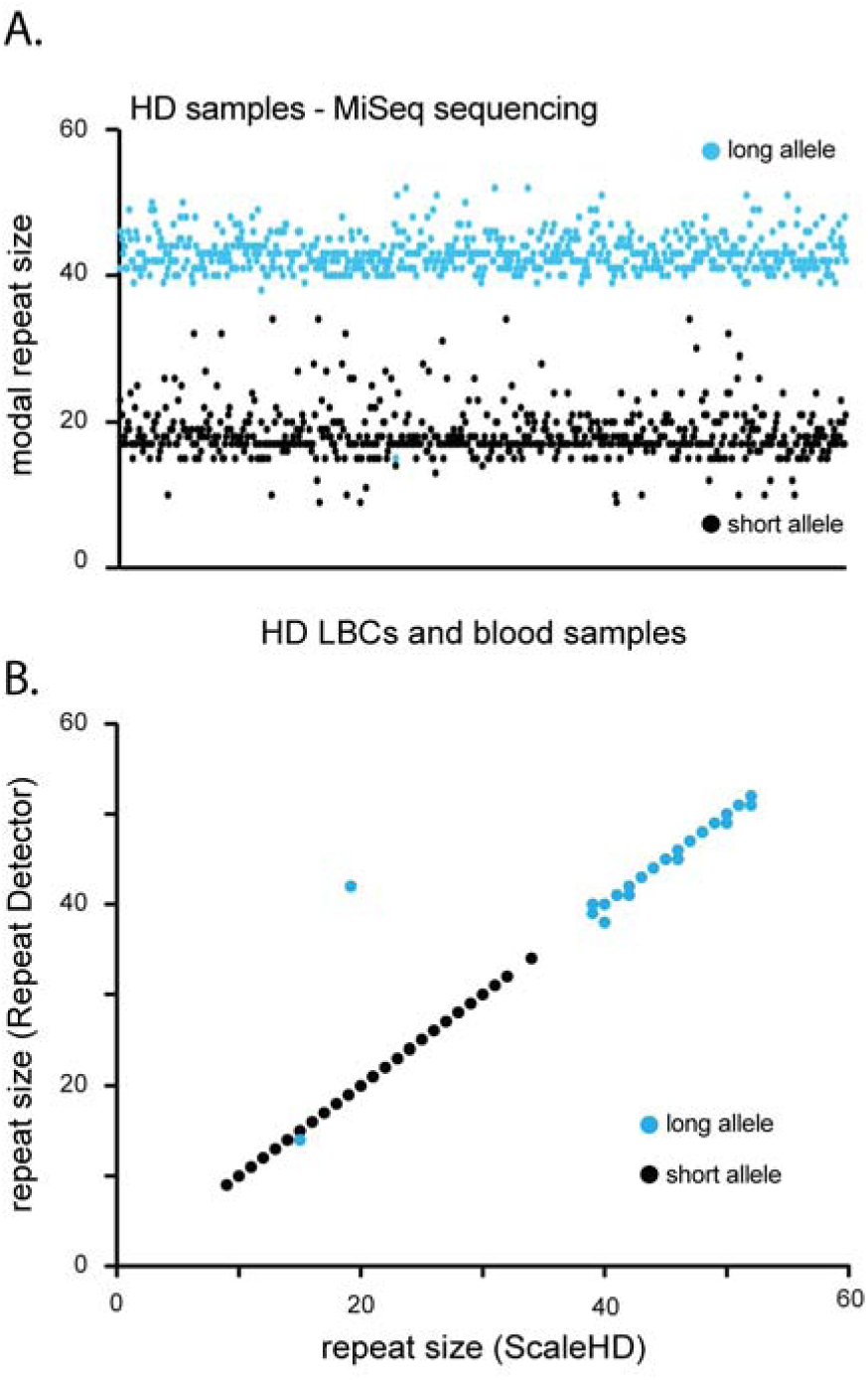
Repeat Detector is highly accurate on HD samples. A) Modal repeat size in the HD MiSeq dataset determined by RD using the restrictive parameters. Each dot is an allele. Blue dots are the longer of the two alleles in a sample, whereas black dots are the shorter alleles.Read depth and mapping metrics for all datasets can be found in Table S4. B) modal repeat size in the HD MiSeq samples comparing ScaleHD and Repeat detector. RD applied to multiple different repeat motifs

### RD is applicable on multiple repeat compositions

To test the applicability of RD to other repeat compositions, we analysed publicly available datasets generated using PCR-free libraries for SMRT (33, 34) and MinION (37, 56) sequencing. These datasets included expanded CAG, CTG, and GGGGCC repeats, as well as short CGG, GGGGCC, and ATTCT repeats. RD found the same repeat size as previously reported for every sample sequenced using SMRT technology (Fig. S5). However, with the MinION sequencing data containing expanded GGGGCC repeats (37), RD dramatically underestimated the repeat size (Fig. S6a). Upon visual inspection of the MinION sequencing reads, we found that the expected repeat motif was too often mutated to be reliably detected (Fig. S6b). This is consistent with Ebert *et al*. (32), who found that when generating whole genome sequences using MinION there was no read aligning to the GGGGCC repeat at the *C9orf72* locus. To determine whether this was indeed due to the quality of MinION sequencing rather than repeat motif composition, we used a recently published MinION dataset that included expanded CAG/CTG repeats from the *HTT* locus (38). We found that only a few sequences were accurate enough to determine repeat size. Most had a very high error rate that prevented us from obtaining accurate repeat counts in this dataset (Fig. S6cd). We conclude that RD is applicable to datasets generated with MinION for this method is too error-prone to identify repeat size down to individual reads.

### RD exposes repeat instability in amplification-free datasets

We next sought to determine whether we would have enough accuracy at the single CCS level to detect heterogeneity of repeat sizes within samples. This was already suggested in the previous datasets with the larger repeat tracts showing more size heterogeneity (Fig. 2). However, in PCR-based library preparation methods there may be slippage errors and other PCR artefacts that may contribute to size heterogeneity, and the distribution of repeat size may not be limited to biological variation (8, 30). Up to 80% of Fuchs endothelial corneal dystrophy (FECD) patients have an expansion of 50 or more CTGs in the third intron of *TCF4* (termed CTG18.1) (57). Here, we analysed a high-quality amplification-free library generated from FECD patient-derived whole blood genomic DNA samples (n=11) displaying a diverse range of CTG18.1 allele lengths and zygosity status (Fig. 4) (33). We found that we could reproduce, for all samples, the modal repeat size determined previously using PacBio’s proprietary RepeatAnalysisTools (Table 1). In addition, repeat instability was obvious with expansion-biased mosaicism, especially for longer alleles (Table 1, Fig. 4 and Fig. S7). We found that RD was largely in agreement with previous studies by Hafford-Tear *et al*. (33) in determining the largest repeat tract present in a sample. In one case, however, RD found a maximum repeat length in one of the samples to be over 1300 units larger than previously identified (566 CTGs identified using RepeatAnalysisTools versus 1875 CTGs with RD, Table 1). Tandem-Genotypes, by contrast, found significantly larger alleles than RD or RepeatAnalysisTools on the expanded alleles, suggesting that it is the more permissive algorithm. Specifically for modal repeat size, it often diverged by a few repeats compared to both RD and RepeatAnalysisTools, with the latter two being in agreement. Together these results show that RD may be used to determine the frequency of repeat instability, in addition to modal repeat size for the FECD SMRT dataset.

**Fig. 4:**
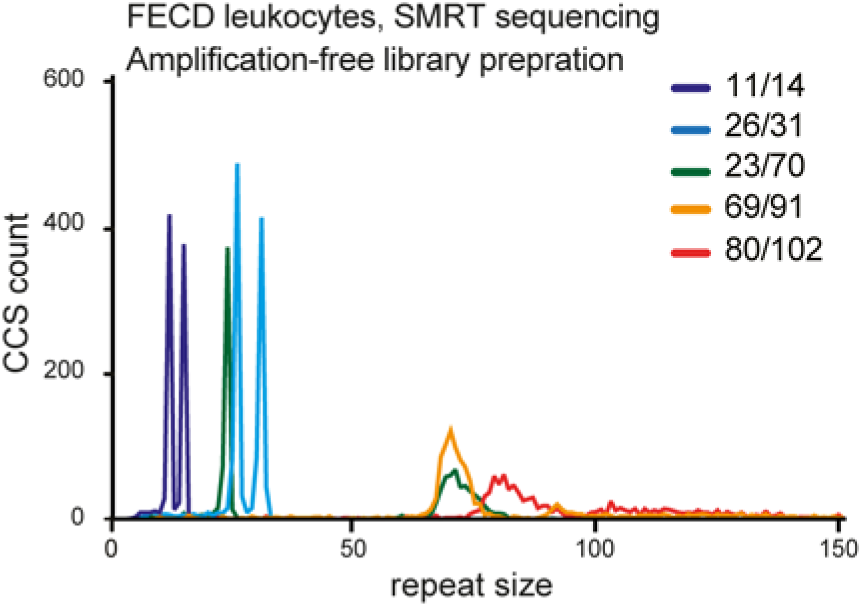
RD exposes repeat instability in amplification-free datasets. Repeat size distribution of the CTG repeat found within a FECD patient cohort from reference (33) prepared using an amplification-free library and SMRT sequenced. Note the wide spread of the repeat sizes on the larger alleles. Only a selection of the samples are plotted for clarity. Density plots for all the samples can be found in Fig. S7. Read depth and mapping metrics for all datasets can be found in Table S4.

**Table 1:** Comparison between RepeatAnalysisTools, Repeat Detector, and Tandem-Genotypes on previously published data for the FECD SMRT dataset.

| Sample | Modal allele Size<br>(no. of repeat units - Short/long alleles) |  |  | Largest Repeat Tract |  |  |
| --- | --- | --- | --- | --- | --- | --- |
|  | RepeatAnalysisTools* | Repeat Detector§ | Tandem-Genotypes | RepeatAnalysisTools* | Repeat Detector§ | Tandem-Genotypes |
| 1 | 11/14 | 11/14 | 11/14 | Not determined | 15 | 18 |
| 2 | 25/30 | 25/30 | 25/30 | 37 | 37 | 37 |
| 3 | 23/70 | 23/70 | 23/68 | 90 | 90 | 90 |
| 4 | 23/73 | 23/73 | 23/74 | 115 | 115 | 116 |
| 5 | 11/80 | 11/80 | 11/81 | 169 | 169 | 170 |
| 6 | 32/110 | 32/110 | 31/110 | 566 | 1875 | 2001 |
| 7 | 17/131 | 9†/131 | 17/126 | 1361 | 1381 | 1393 |
| 8 | 80/102 | 80/102 | 80/102 | 498 | 498 | 506 |
| 9 | 72/118 | 72/118 | 73/117 | 1593 | 1285 | 2221 |
| 10 | 69/91 | 69/91 | 69/91 | 1014 | 1047 | 1050 |
| 11 | 79/141 | 79/141 | 78/140 | Not determined | 1581 | 1580 |

\*Data from (33)
§The restrictive parameters were used to determined repeat size.
†This lower number is due to the presence of an interruption in this allele. This is evident when the permissive parameters are also used (see Fig. 5).

### Identifying interrupted alleles using RD

When optimising RD, we settled on two sets of parameters, one that allowed for the occurrence of sequencing errors (permissive) and one that did not (restrictive). The analyses presented above were conducted using the restrictive parameters. On the HD MiSeq dataset, the restrictive parameters returned the length of the pure repeat tract whereas the permissive parameters count the downstream interruption and the first triplet downstream of the repeat tract, typically CCG. Thus, alleles with the canonical CAACAG interruption will yield a difference of 3 repeats between the permissive and restrictive parameters (Fig. 5ab). By contrast, alleles without the interruption yield only one repeat difference between the two profiles (Fig. 5ab) and the ones with a duplicated CAACAG motif show a difference of 5 units (Fig. 5ac). The shifts can be used to identify samples with repeat interruptions or unusual allele structures and narrow down which samples need to be inspected manually. Using this approach, we could accurately identify the sole sample in the HD MiSeq dataset with a CAC interruption within its CAG repeat (Fig. 5d) and the interrupted non-pathogenic allele in Sample 7 of the FECD dataset (Fig. 5e). We could also identify a previously unknown 111bp insertion in the GFP(CAG)_308_ cell line (Fig. 5f), which we confirmed by PCR and Sanger sequencing, as well as in a separate flowcell. These results suggest that RD can be used to identify individual alleles with interruptions at multiple different loci.

**Fig. 5:**
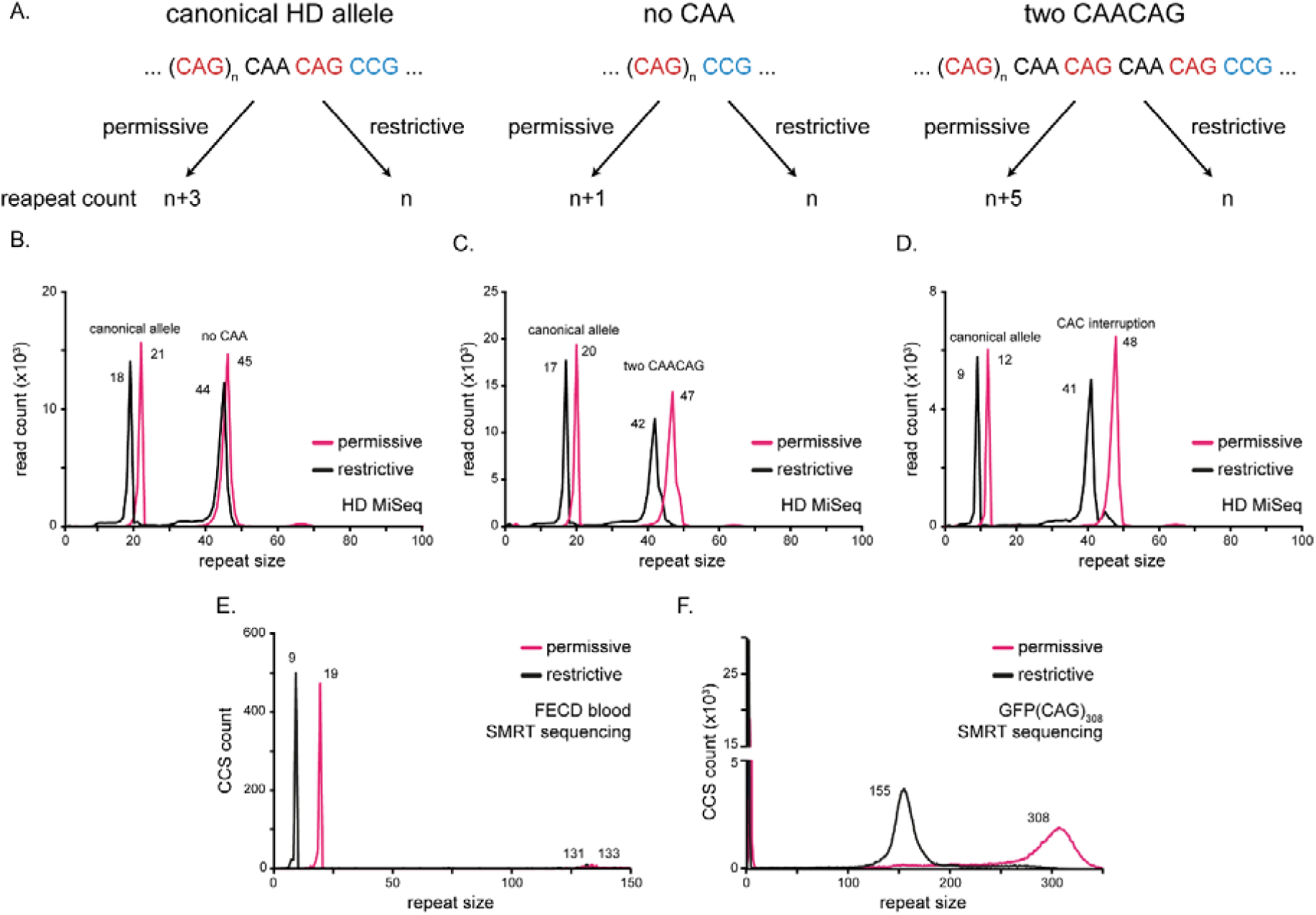
Identifying samples with interruptions using RD. A) Interruptions at the 3’ end of HD alleles can be distinguished using the difference in repeat size between RD’s permissive and restrictive parameters. For instance, the most common allele (left), containing a CAA interruption will return a difference of 3 repeats between the parameter settings. By contrast, an allele without the CAA (middle) or with two CAACAG motifs return differences of 1 and 5, respectively. B) Example of a sample from the HD MiSeq dataset with a canonical non-pathogenic allele, and an expanded allele without a CAA interruption. C) Example for a HD MiSeq sample with a canonical short allele and an expanded allele with a duplicated CAACAG motif. D) One of the samples contained a rare CAC interruption in the repeat tract that returns a difference larger than expected from the known alleles. E) A previously known interrupted allele in a FECD sample (33) was correctly identified. F) Our GFP(CAG)308 line was found to have an insertion of 111bp after 155 CAG repeats.

## Discussion

Here we developed and applied RD, which detects and counts tandem repeats in targeted sequencing data. RD was as accurate as ScaleHD on the HD MiSeq dataset and as Tandem-Genotypes and RepeatAnalysisTools on the FECD SMRT dataset. RD could also identify interruptions, when present, as readily as RepeatAnalysisTools on the FECD SMRT dataset. None of the other available algorithms could be used with all of these datasets. For example, ScaleHD can identify known interruptions only at the *HTT* locus by adding them to its library of sequences whereas RepeatAnalysisTools can only be applied to amplification-free SMRT sequencing. Tandem-Genotypes could also be applied to multiple loci, but it is not designed to find interruptions. Tandem-Genotypes also requires a specific aligner, LAST (49), which does not work with artificial constructs such as our GFP reporter. Thus, the main strength of RD is its versatility: it works on multiple different sequencing platforms, multiple loci, including artificial reporters, and can identify interrupted alleles readily. Although RD allows for changing parameter scores to accommodate the systematic sequencing errors of each sequencing platform, we did not have to change the parameters when applying it to SMRT and MiSeq, or when we applied it to different loci or repeat compositions. Further optimisation of the weighting profiles may help to compensate for the higher error rate of MinION sequencing datasets.. RD could detect repeat instability in HD and FECD blood-derived samples prepared with a PCR-based or amplification-free protocol, respectively. In the amplification-free TCF4 PacBio dataset where PCR biases against the longer repeats could be ruled out, some samples had large expansions with some reads having several hundreds of repeats. This is not uncommon in FECD patient-derived samples, but they are difficult to detect by any method, except perhaps for small-pool PCR followed by Southern blotting (58). Our data, together with that of a recent pre-print on DM1 (29), suggest that it is possible to detect repeat instability as well as interruptions in PCR-free sequencing methods. More work needs to be done to validate this approach. Specifically, comparing samples with different levels of repeat instability using both small-pool PCR and amplification-free SMRT libraries will be critical. Notably, RD would not be suitable for whole genome sequencing datasets and these datasets would not be suitable to determine repeat size mosaicism.

Several datasets used Oxford Nanopore sequencing on expanded repeats (32, 37–39), yet levels of repeat mosaicism was only reported in one study (39). This is likely because the error rate of MinION is too high to be confident about the size of the repeats in individual reads. On non-repetitive loci, this is not a problem because sequencing with a high coverage can compensate for stochastic errors in individual reads. On an unstable tandem repeat, however, this averages out the repeat size differences between reads and the distribution of the repeat size is lost. Oxford Nanopore is currently too error-prone for use to determine repeat size heterogeneity within a sample it can only be used to obtain modal repeat size. Improvements to base calling may help mitigate this issue.

Current sequencing efforts have been limited to modal repeat sizes below about 150 CAGs, with the notable exceptions of myotonic dystrophy samples (23, 29). Here we could detect repeat sizes in excess of 1800 CTGs at the *TCF4* locus in individual reads. It will be interesting to test how well RD performs on datasets with longer repeats as those become available.

Interruptions within the repeat tract are classically detected using repeat-primed PCR, whereby a primer sits in the flanking sequence and another within the repeat tract itself (23, 59). This leads to a pattern on capillary electrophoresis with a periodicity the size of the repeated unit and of decaying intensity. Interruptions appear in the intensity traces as gaps in places where the repeat primer could not bind. Depending on the position of the interruption within the repeat tract, these may be difficult to detect accurately, especially if they are far from the 3’ or 5’ ends of the repeat tracts. Once an interruption is detected, its identity and position need to be confirmed by Sanger sequencing or restriction digest. Targeted sequencing coupled with RD would identify first the presence of an interruption in the sample, and then the examination of individual reads would reveal both the position and the content of the interruption. This would dramatically speed up the process and may thereby reduce cost.

In its current version, RD has a few limitations. One is that it requires user intervention to identify the nature of the interruption detected in a sample and cannot discriminate between single and multiple interruptions in the same allele. This will be important to address as several alleles from DM1 patients, for example, with complex interruptions have been documented (23, 29). In these samples, RD would return the size of the longest interruption-free repeat stretch. Moreover, the size of the interruption tolerated by the permissive parameters depends on the position of the interruption and on the number of repeated units flanking the insertion. For example, the larger interruption found in the GFP(CAG)_308_ line was allowed with the permissive parameters because it was flanked by two repeat tracts of 155 and 115 repeats. Thus, in some cases, large interruptions may not be found, or the parameters may need to be adjusted. This was highlighted by the Oxford Nanopore datasets that we analysed here. RD ignores flanking sequences and thus would be blind to, for example, the significant polymorphism found in the CCG repeat downstream of the HD allele (8). To get around this, RD could be run once for the size of the CAG repeat and once for the size of the CCG repeats and its interruptions downstream of the repeat tract. Improvements to RD may also include changes to the weighting scores for improved accuracy on MinION datasets and on a wider variety of repetitive sequences (e.g., telomeres).

Some tandem repeats may not benefit from RD. For example, Variable Number Tandem Repeats (VNTRs) are not pure and often contain multiple different repeated motifs. In these cases, we would expect RD to be able to count the repeats provided that the permissive weighting scores are adjusted. The restrictive parameters would then return the longest stretch of pure repeats. Thus, for highly interrupted repeats RD would perform similarly as on error-riddled reads.

We have shown that RD can accurately determine repeat size from targeted sequencing data from SMRT, MiSeq, and MinION sequencing platforms. It is not limited by a requirement for a library of reference sequences, can be applied to a wide variety of disease loci and repeat compositions, can be used to identify alleles with interruptions, and can document repeat length mosaicism within a sample. Together, these characteristics make RD broadly applicable and capable tool for analysis of expanded tandem repeats.

## Acknowledgements

We thank John H. Wilson, Alvaro Murillo Bartolome, Helder Ferreira, Fisun Hamaratoglu, and Andrew Seeber for comments on the manuscript. Mark Ebbert kindly provided the Nanopore MinION data from ALS individuals. The Registry HD lymphoblastoid DNA and cell lines were provided by the European Huntington’s Disease Network project #984. The Registry study is supported by the European Huntington’s Disease Network (EHDN), funded by CHDI Foundation, Inc. The funding source had no role in study design; in the collection, analysis, and interpretation of data; in the writing of the report; and in the decision to submit the paper for publication. We acknowledge the support of the Supercomputing Wales project, which is part-funded by the European Regional Development Fund (ERDF) via the Welsh Government.

## Conflict of interests

LJ is on the scientific advisory boards of LoQus23 Therapeutics and Triplet Therapeutics and a member of the Executive Committee of the European Huntington’s Disease Network. Within the last five years DGM has been a scientific consultant and/or received an honoraria/stock options/research contracts from AMO Pharma, Charles River, LoQus23, Small Molecule RNA, Triplet Therapeutics, and Vertex Pharmaceuticals. Within the last 5 years AED has been a scientific consultant for and/or received an honoraria/stock options/research contracts from Triplet Therapeutics, LoQus23 Therapeutics, Design Therapeutics, ProQR Therapeutics and Prime Medicine.

## Data availability

The source code and dependencies are available here: https://github.com/DionLab/RepeatDetector. The HD SMRT and GFP(CAG)_x_ SMRT datasets are available from the Gene Expression Omnibus (GSE199005). The GFP(CAG)_x_ lines are available upon request from the corresponding author.

## Author contributions

AST analysed the interruptions and repeat size in the MiSeq HD, SMRT sequencing HD, and in the FECD SMRT sequencing datasets. She generated the tables and figures together with VD, DB, and EH. DB optimised RD for CCS, analysed the repeat size on the HD and GFP SMRT sequencing datasets, and initiated the analysis of repeat size in the HD MiSeq dataset. NG tested the beta version of RD and optimised the parameters used here. TS created and programmed RD under the supervision of IX. LA, ER, ET, and PB optimised and produced PacBio sequencing libraries. JM advised on library preparation, trained ER, ran the sequencing, and generated the CCSs. EH analysed the GFP(CAG)_x_ SMRT datasets. BM generated the MiSeq sequences together with MC under the supervision of DGM, TM, and LJ. NJHT provided raw data for the FECD PCR-free dataset under the supervision of AED. VD, AST, DB, NG, and IX designed the experiments. VD wrote the manuscript and all the co-authors provided input and feedback.

## Funding

The Dion lab is funded by a Professorship from the Academy of Medical Sciences (AMSPR1\1014) and by the UK Dementia Research Institute, which receives its funding from DRI Ltd, funded by the UK Medical Research Council, Alzheimer’s Society and Alzheimer’s Research UK. BM received a Cardiff University School of Medicine Studentship. LJ was supported by an MRC Centre grant (MR/L010305/1); DGM, MC, LJ and THM by CHDI. THM was supported by a Welsh Clinical Academic Track Fellowship, an MRC Clinical Training Fellowship (MR/P001629/1) and a Patrick Berthoud Charitable Trust Fellowship through the Association of British Neurologists. BM, TM and LJ were supported by the Brain Research Trust (201617-06).

## Notes

https://github.com/DionLab/RepeatDetector

https://github.com/DionLab/libprf

### Competing Interest Statement

LJ was on the scientific advisory boards of LoQus23 Therapeutics and Triplet Therapeutics and a member of the Executive Committee of the European Huntington's Disease Network. Within the last five years DGM has been a scientific consultant and/or received an honoraria/stock options/research contracts from AMO Pharma, Charles River, LoQus23, Small Molecule RNA, Triplet Therapeutics, and Vertex Pharmaceuticals. Within the last 5 years AED has been a scientific consultant for and/or received an honoraria/stock options/research contracts from Triplet Therapeutics, LoQus23 Therapeutics, Design Therapeutics, ProQR Therapeutics and Prime Medicine.

### Summary of Updates

This version of the manuscript was updated to improve much of the text. The discussion has been extended. It includes further benchmarking using Tandem-Genotype. A figure and a figure panel in the supplementary material were added to provide further details on the parameters used for RD.

