## Supplementary material for "Repeat Detector: versatile sizing of expanded tandem repeats and identification of interrupted alleles from targeted DNA sequencing": Suppl

1. UK Dementia Research Institute at Cardiff University, Hadyn Ellis Building, Maindy Road, Cardiff, CF24 4HQ, UK
2. Centre for Integrative Genomics, University of Lausanne, Bâtiment Génopode, 1015 Lausanne, Switzerland
3. Vital-IT Group, Swiss Institute of Bioinformatics, 1015 Lausanne, Switzerland
4. Current address: Newbiologix, Ch. De la corniche 6-8, 1066 Epalinges, Switzerland
5. MRC Centre for Neuropsychiatric Genetics and Genomics, Cardiff University, Hadyn Ellis Building, Maindy Road, Cardiff CF24 4HQ, UK
6. Current address: Molecular Neurogenetics Unit, Center for Genomic Medicine, Massachusetts General Hospital, Boston, MA 02114, USA.
7. Current address: Sorbonne Université, École normale supérieure, PSL University, CNRS, Laboratoire des biomolécules, LBM, 75005 Paris, France.
8. Institute of Molecular, Cell and Systems Biology, College of Medical, Veterinary and Life Sciences, Davidson Building, University of Glasgow, Glasgow, G12 8QQ, UK
9. UCL Institute of Ophthalmology, 11-43 Bath Street, London, EC1V 9EL UK
10. http://www.ehdn.org/wp-content/uploads/REGISTRY-contributors-full-list.pdf
11. Health2030 Genome Center, Ch des Mines 14, 1202 Genève, Switzerland

*: These authors contributed equally to this work.

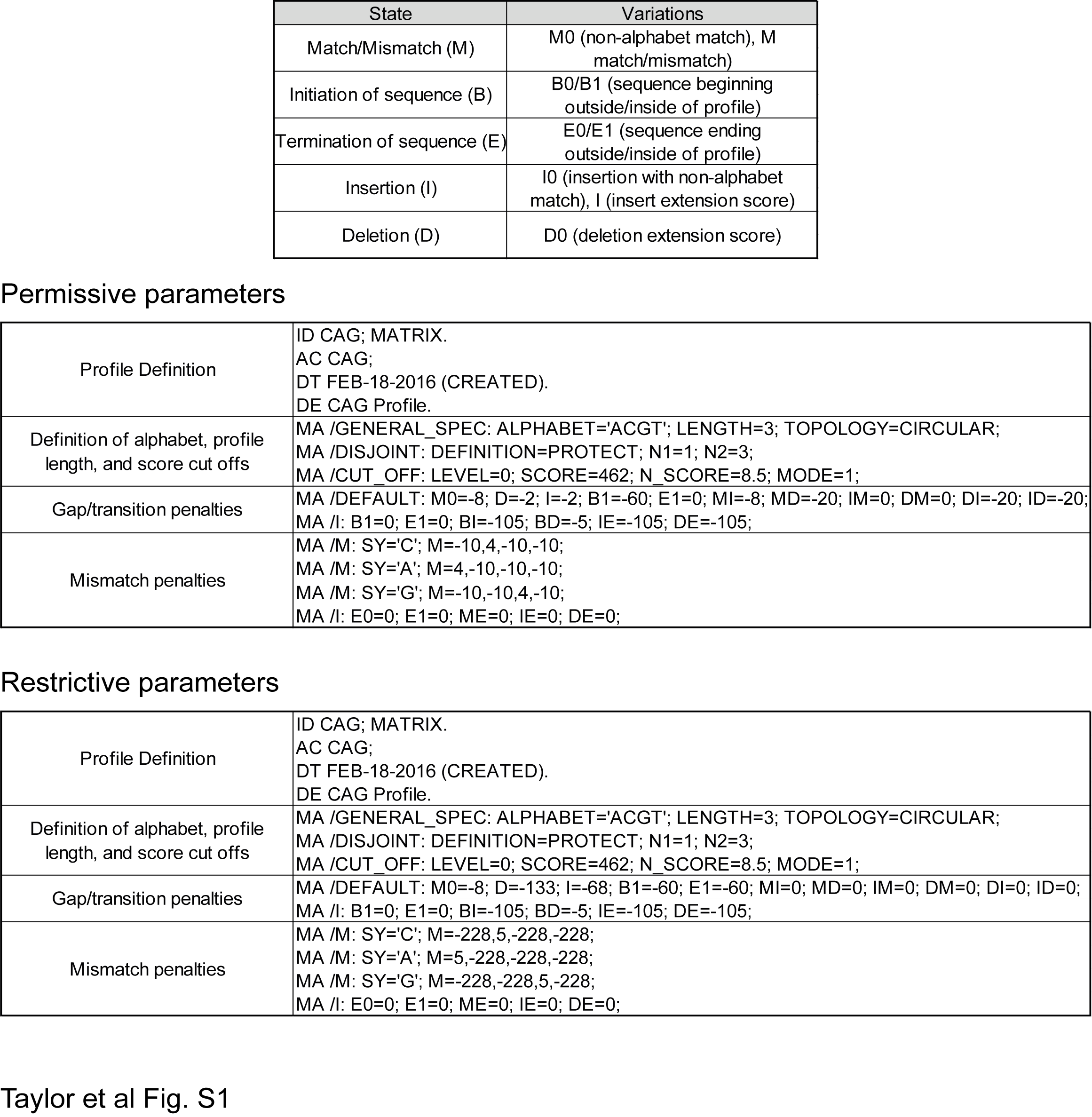

***Fig. S1:*** *Weighting scores and parameters for the restrictive and permissive profiles used in RD.*

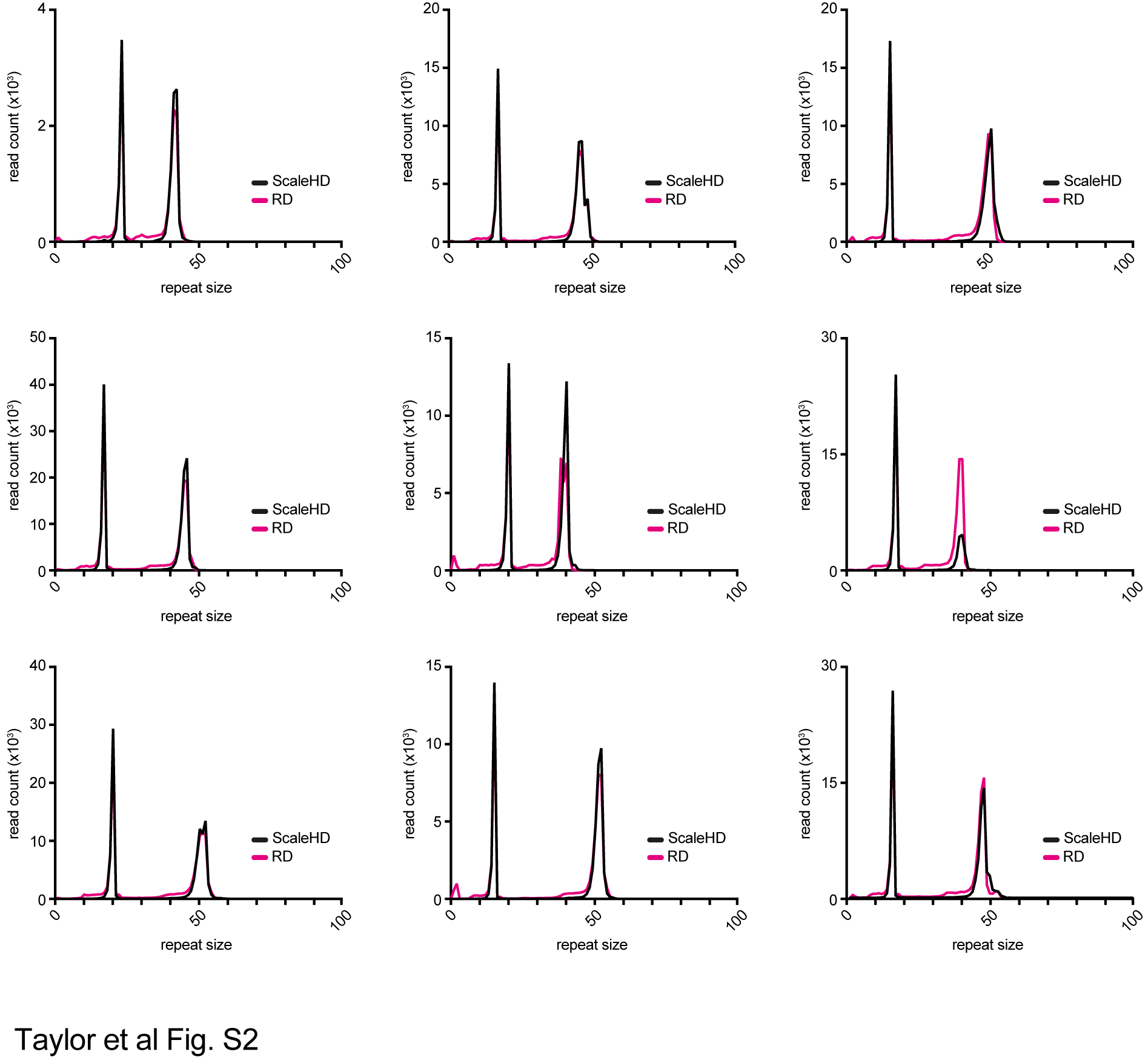

***Fig. S2:*** *Comparison of RD and ScaleHD on 9 samples that differed by one or two repeats. For RD, we used the restrictive parameters.*

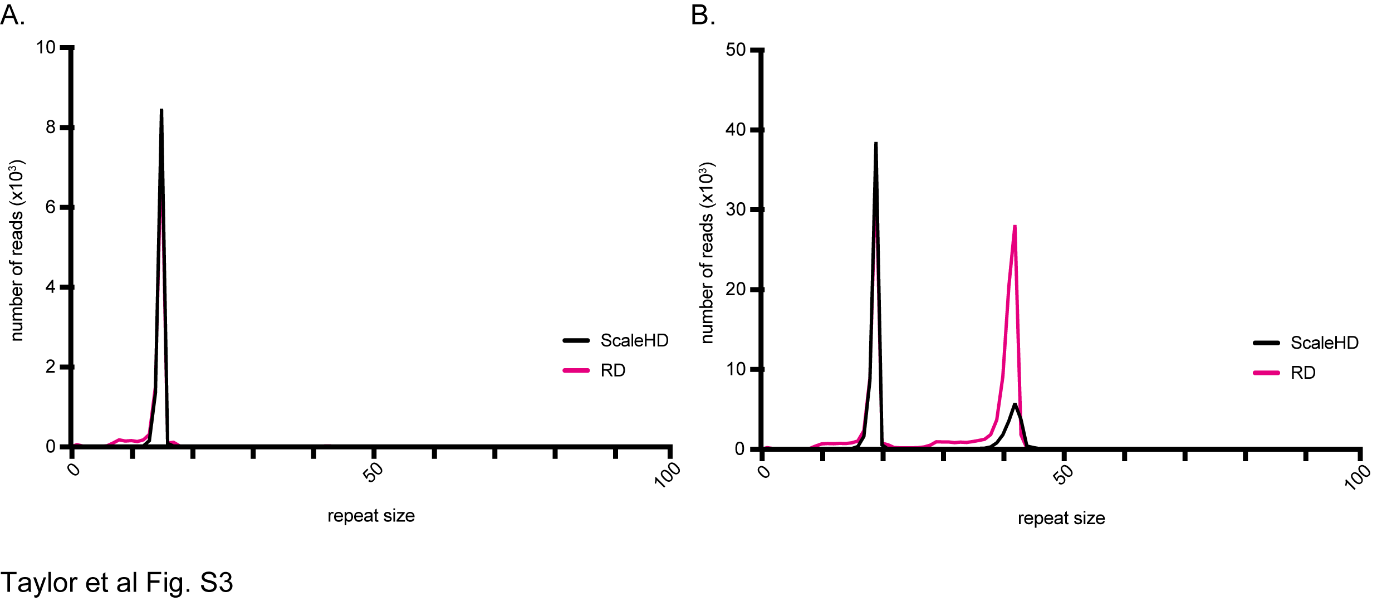
***Fig. S3:*** *Density plot of samples that differed in their modal repeat count between ScaleHD (black) and Repeat Detector (restrictive parameters - pink). A) This sample is a homozygous that ScaleHD called correctly. RD used the two most common allele sizes and thus returned a heterozygous sample with one repeat difference. B) In this sample, ScaleHD discarded many reads that did not align well to the library of sequences for undefined reasons. Other samples from the same individual did not have that issue. RD, not needing this alignment step to a reference sequence beyond the repetitive motif, called both alleles correctly.*

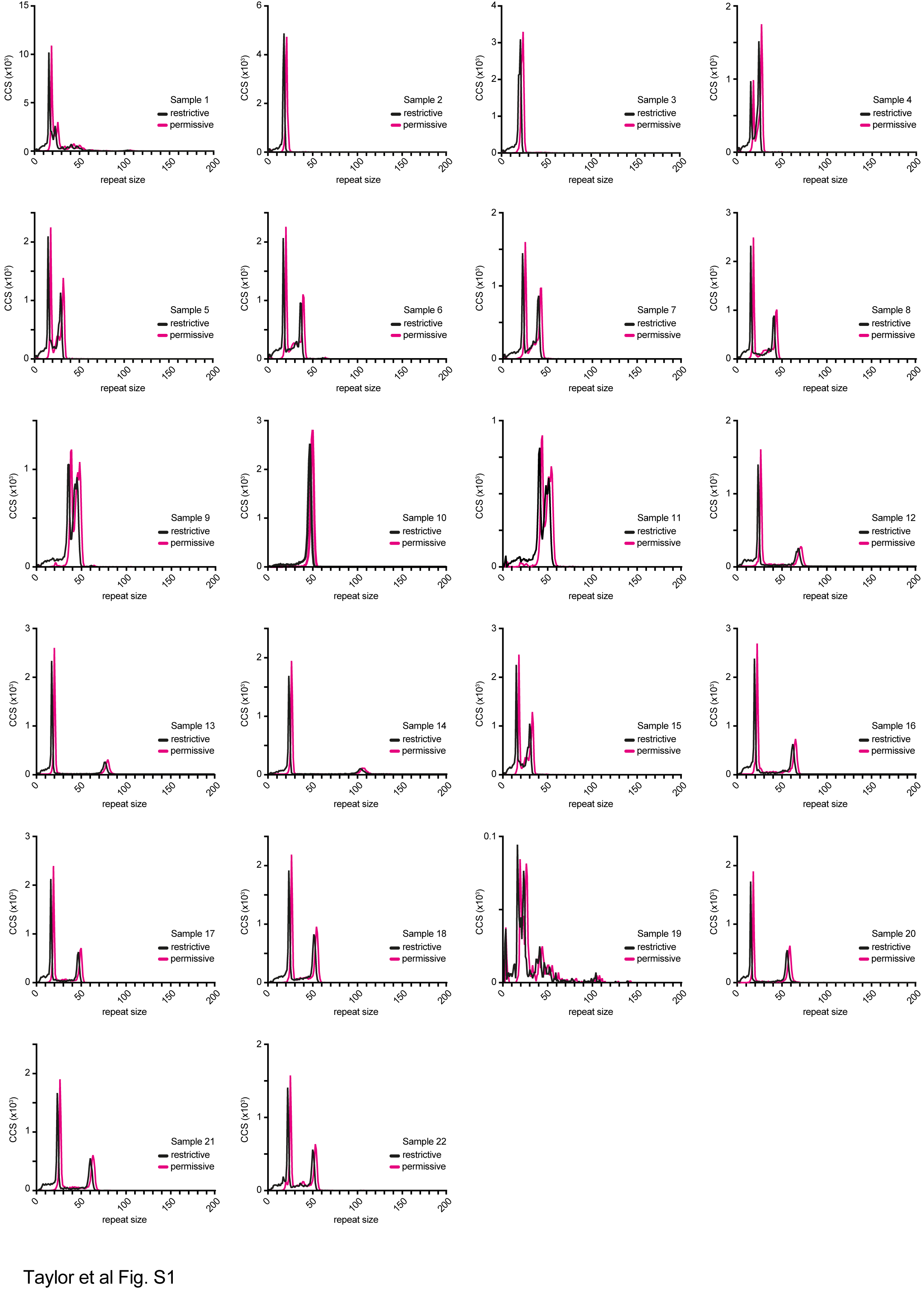

***Fig. S4:*** *RD density plots for all the samples in the HD SMRT dataset. Sample information can be found in Table S1.*

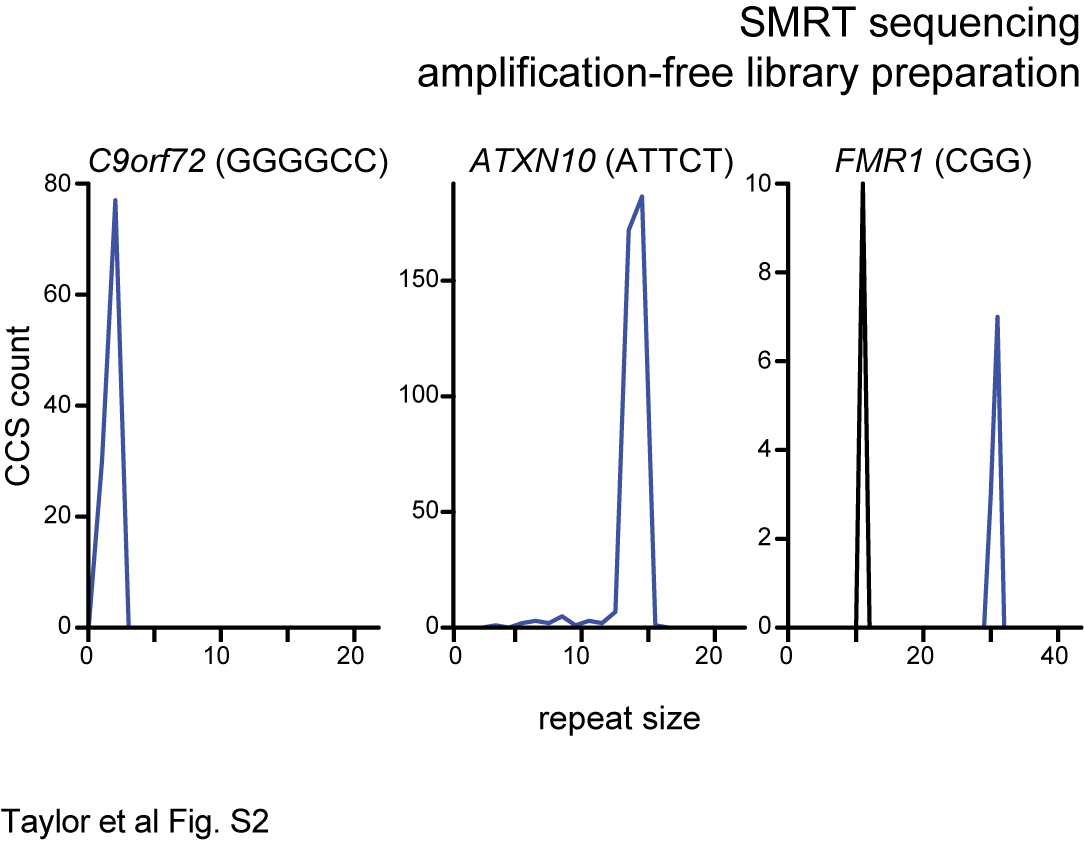

***Fig. S5:*** *Representative RD density plots for the non-pathogenic alleles at three non-CAG loci in an HD individual. The sequencing data is from* (1)*.*

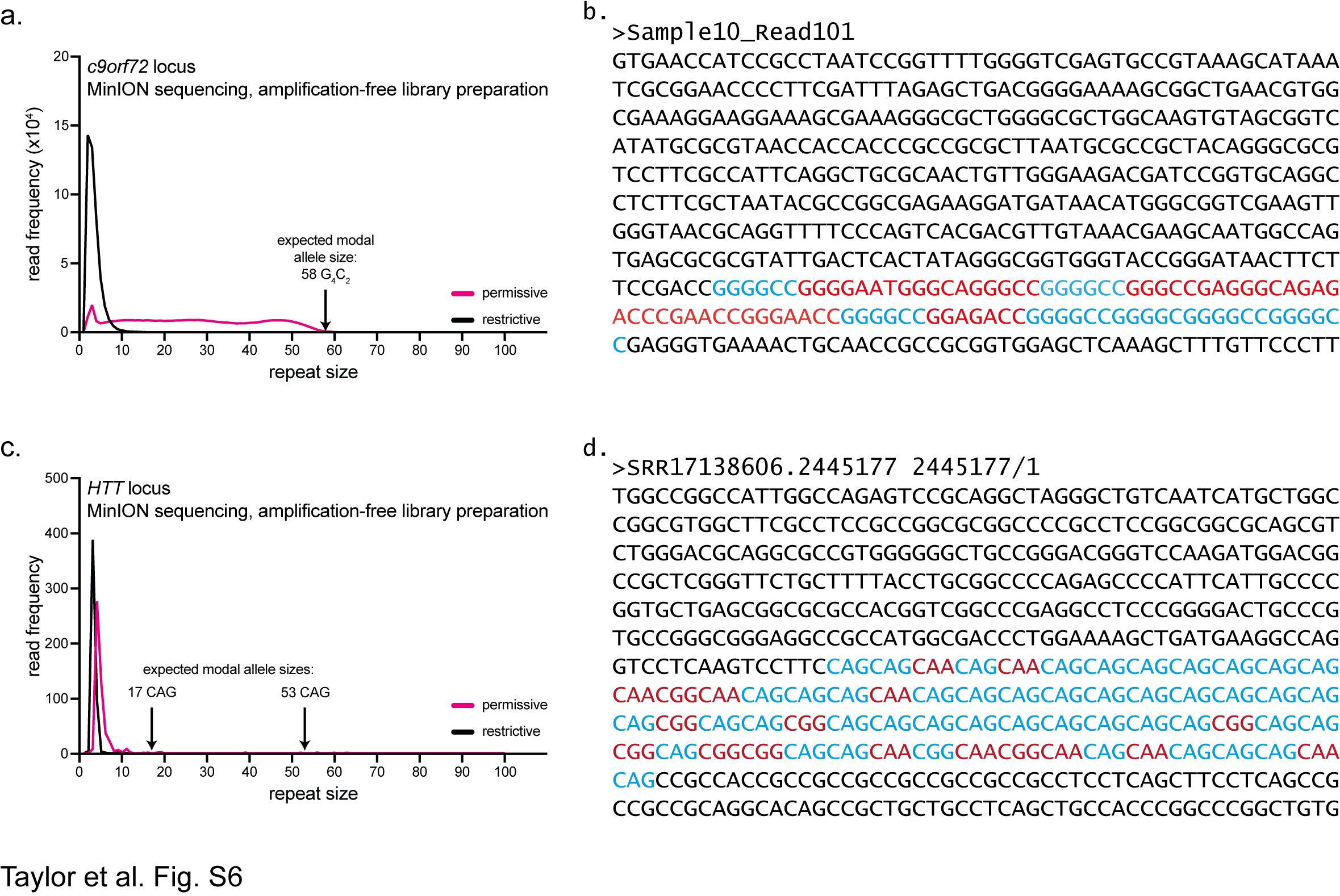

***Fig. S6:*** *RD returns short repeat sizes on MinION sequencing data due to the high error-rate. A) Repeat size distribution analysed using RD’s restrictive and permissive parameters for the* C9ORF72 *locus from plasmid template (Sample 10 from* (2)*). Arrow is expected repeat size. B) A representative read with the expected G_4_C_2_ motif in blue and sequencing errors in red. C) Density plot of repeat size distribution at* HTT *in sample NA13507 from MinION sequencing data published in* (3)*. D) A representative read with the expected CAG motif in blue and sequencing errors in red.*

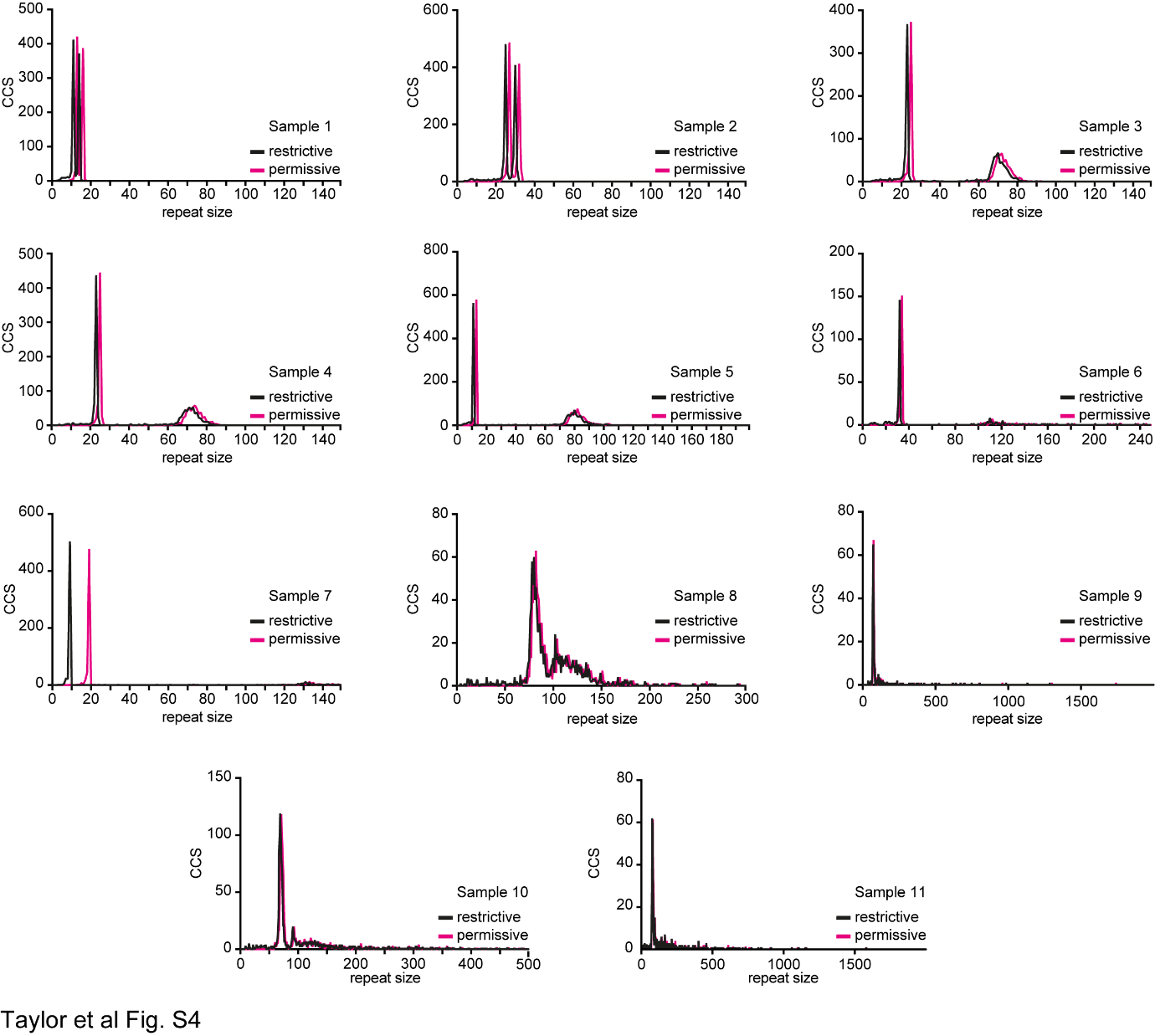

***Fig. S7:***  *RD density plots of the FECD SMRT dataset. Sample numbering is the same as in Hafford-Tear et al* (4)*.*

**Table S1:** HD SMRT sequencing samples, barcodes and primers used.

| **Flowcell** | **Sample*** | **Barcode (on forward primer)** | **Forward primer** | **Reverse primer** | **Forward Primer name** | **Reverse primer name** |
| --- | --- | --- | --- | --- | --- | --- |
| 1 | NA20245 | TCAGACGATGCGTCAT | GCCTCCCTTACCATGCAGT | GCCCTCTTCCCTCTCAGACT | oVIN2492 | oVIN1566 |
| 1 | NA20206 | CTATACATGACTCTGC |  |  | oVIN2493 |  |
| 1 | NA20207 | TACTAGAGTAGCACTC |  |  | oVIN2494 |  |
| 1 | NA20246 | TGTGTATCAGTACATG |  |  | oVIN2495 |  |
| 1 | NA20247 | ACACGCATGACACACT |  |  | oVIN2496 |  |
| 1 | NA20248 | GATCTCTACTATATGC |  |  | oVIN2497 |  |
| 1 | NA20249 | ACAGTCTATACTGCTG |  |  | oVIN2498 |  |
| 1 | NA20250 | ATGATGTGCTACATCT |  |  | oVIN2499 |  |
| 1 | NA20208 | CTGCGTGCTCTACGAC |  |  | oVIN2500 |  |
| 1 | NA20209 | GCGCGATACGATGACT |  |  | oVIN2501 |  |
| 1 | NA20251 | CGCGCTCAGCTGATCG |  |  | oVIN2502 |  |
| 1 | NA20252 | GCGCACGCACTACAGA |  |  | oVIN2503 |  |
| 1 | NA20210 | ACACTGACGTCGCGAC |  |  | oVIN2504 |  |
| 1 | NA20253 | CGTCTATATACGTATA |  |  | oVIN2505 |  |
| 1 | GM02168 | ATAGAGACTCAGAGCT |  |  | oVIN2506 |  |
| 1 | GM03620 | TAGATGCGAGAGTAGA |  |  | oVIN2507 |  |
| 1 | GM13504 | CATAGCGACTATCGTG |  |  | oVIN2508 |  |
| 1 | GM13505 | CATCACTACGCTAGAT |  |  | oVIN2509 |  |
| 1 | GM13506 | CGCATCTGTGCATGCA |  |  | oVIN2510 |  |
| 1 | GM13507 | TATGTGATCGTCTCTC |  |  | oVIN2511 |  |
| 1 | GM13508 | GTACACGCTGTGACTA |  |  | oVIN2512 |  |
| 1 | GM14044 | CGTGTCGCGCATATCT |  |  | oVIN2513 |  |
| 2 | GFP(CAG)_15_ | CACACGCGCGTGCTCG | AAGAGCTTCCCTTTACACAACG | TACTTGTACAGCTCGTCCATGC | oVIN0881 | oVIN2514 |
| 2 | GFP(CAG)_51_ | TCTCTCACAGTCGAGC |  |  | oVIN0887 |  |
| 2 | GFP(CAG)_91_ | CGCGCGTGTGTGCGTG |  |  | oVIN0879 |  |
| 2 | GFP(CAG)_308_ | GTCATCACACATCTCT |  |  | oVIN0889 |  |
| 3 | GFP(CAG)_15_ | CACACGCGCGTGCTCG |  |  | oVIN0881 |  |
| 4 | GFP(CAG)_51_ | TCTCTCACAGTCGAGC |  |  | oVIN0887 |  |
| 5 | GFP(CAG)_91_ | CGCGCGTGTGTGCGTG |  |  | oVIN0879 |  |
| 6 | GFP(CAG)_308_ | GTCATCACACATCTCT |  |  | oVIN0889 |  |

*: Samples starting with NA were DNA samples obtained directly from Coriell. GM samples were LBCs obtained from Coriell and grown in our laboratory prior to DNA isolation.

**Table S2:** Primers used to sequence the repeat tract in GFP(CAG)_x_ cells and to verify the presence of the duplication in GFP(CAG)_308_.

| **Primer** | **Sequence** | **Reference** |
| --- | --- | --- |
| oVIN-459 | AAGAGCTTCCCTTTACACAACG | (5) |
| oVIN-460 | TCTGCAAATTCAGTGATGC | (5) |

**Table S3:** Selected samples for sequencing (total N=652).

| **DNA type** | **Number sequenced*** | **Number successfully sequenced** |
| --- | --- | --- |
| LBCs (Early samples) | 250 | 249 |
| LBCs (Late samples) | 250 | 248 |
| Time course LBCs | 47 | 47 |
| Blood DNA | 98 (49) | 98 |
| Positive controls | 7 (1) | 7 |

*Numbers in brackets indicate the actual number of individuals for each group; blood DNA were all sequenced twice, and the single positive control was sequenced on each plate for a total of 7 times. Blood DNA consisted of the 7 expected onset individuals as well as 26 early and 16 late.

**Table S4:** Sequencing metrics for all the datasets analysed

See TableS4_SequencingMetrics.xlsx attached.
